# Three-dimensional culture in a bioengineered matrix and somatic cell complementation to improve growth and survival of bovine preantral follicles

**DOI:** 10.1101/2024.07.18.604061

**Authors:** Juliana I. Candelaria, Ramon C. Botigelli, Carly Guiltinan, Ariella Shikanov, Anna C. Denicol

## Abstract

**Purpose:** Here we explored poly(ethylene glycol) (PEG) bioengineered hydrogels for bovine preantral follicle culture with or without ovarian cell co-culture and examined the potential for differentiation of bovine embryonic stem cells (bESCs) towards gonadal somatic cells to develop a system more similar to the ovarian microenvironment.

**Methods:** Bovine preantral follicles were first cultured in two-dimensional (2D) control or within PEG hydrogels (3D) and then co-cultured within PEG hydrogels with bovine ovarian cells (BOCs) to determine growth and viability. Finally, we tested conditions to drive differentiation of bESCs towards the intermediate mesoderm and bipotential gonad fate.

**Results:** Primary follicles grew over the 10-day culture period in PEG hydrogels compared to 2D control. Early secondary follicles maintained a similar diameter within the PEG while control follicles decreased in size. Follicles lost viability after co-encapsulation with BOCs; BOCs lost stromal cell signature over the culture period within hydrogels. Induction of bESCs towards gonadal somatic fate under WNT signaling was sufficient to upregulate intermediate mesoderm (*LHX1*) and early coelomic epithelium/bipotential gonad markers (*OSR1*, *GATA4*, *WT1*). Higher BMP4 concentrations upregulated the lateral plate mesoderm marker *FOXF1*. *PAX3* expression was not induced, indicating absence of the paraxial mesoderm lineage.

**Conclusions:** Culture of primary stage preantral follicles in PEG hydrogels promoted growth compared to controls; BOCs did not maintain identity in the PEG hydrogels. Collectively, we demonstrate that PEG hydrogels can be a potential culture system for early preantral follicles pending refinements, which could include addition of ESC-derived ovarian somatic cells using the protocol described here.

**CAPSULE SUMMARY:** We demonstrate that three-dimensional bioengineered hydrogels could aid in the survival and growth of small bovine preantral follicles. Moreover, bovine embryonic stem cells have the potential to differentiate towards precursors of somatic gonadal cell types, presenting an alternative cell source for preantral follicle co-culture.

## INTRODUCTION

Folliculogenesis is a complex and selective process resulting in production of a limited quantity of meiotically and developmentally competent oocytes over a female’s lifespan. The molecular processes that coordinate the activation, growth and maturation of ovarian follicles have yet to be fully elucidated, especially during the early preantral stages of development. Progress in understanding preantral folliculogenesis in non-rodent species has been hindered by the lack of appropriate in vitro culture systems and an insufficient knowledge base of ovarian development and function. This is particularly true in large mammal (i.e. cows and humans) preantral folliculogenesis which have prolonged duration and sizeable volumetric changes between the primordial and pre-ovulatory follicle stages. A great advantage of performing in vitro folliculogenesis beginning at the preantral stage includes establishing an alternative fertility preservation and restoration method by utilizing the vast preantral follicle pool for oocyte production [1]. More specifically, this approach would be beneficial for prepubertal girls suffering from blood-borne malignancies as isolated preantral follicles could be matured in vitro without the risk of reintroducing malignant cells or surgical intervention, as could be the case with the culture and reintroduction of ovarian tissue fragments.

Recapitulation of bovine preantral folliculogenesis has been attempted by several groups, but there has been little success in efficiently growing follicles to the preovulatory stage and yielding metaphase II oocytes. Although 2-dimensional culture systems have shown some promise in supporting bovine preantral follicle growth [2–4], mounting evidence suggests that 3-dimensional (3D) culture systems better promote development due to their ability to maintain a follicle’s spherical configuration and preserve cell contact both within the follicular unit (i.e. granulosa cell and oocyte interface) and outside the basement membrane where theca cells are located [5–8]. Matrigel and alginate are commonly used biomaterials to create hydrogels for in vitro follicle culture, however pitfalls include uncontrollable degradation and biological inertness, respectively [9]. Therefore, a highlighted alternative is the use of bioengineered 3D hydrogel culture systems that permit timely degradation by the follicle yet retaining the mechanical support necessary to keep its spherical structure [10,11]. Poly(ethylene glycol) (PEG) hydrogels constructed with proteolytically-degradable peptide crosslinkers have been employed for mouse in vitro folliculogenesis and demonstrated success in growing preantral follicles to the antral stage and producing metaphase II oocytes [12]. Likewise, the addition of adipose-derived stem cells or conditioned media to PEG hydrogels containing mouse preantral follicles further aided development and survival [13]. Despite the promise demonstrated in mice, the use of PEG hydrogels for large mammal in vitro preantral follicle culture has not been reported. Moreover, investigating cell supplementation in the hydrogel system to determine their propensity to become theca cells and/or advantageous effect on follicle growth has not been directly assessed. Indeed, early preantral follicles do not contain theca cells and must recruit precursor cells from the stromal environment to build the essential theca layer, hence addition of cells to the system would be essential when starting with primordial- or primary-stage follicles. More specifically, the hedgehog signaling pathway is key to driving theca cell differentiation and recruitment [14–16], therefore cells added to the culture system should harbor the cellular machinery to respond to hedgehog ligands secreted by the follicular granulosa cells.

Although native ovarian stromal cells have been used as feeder cells for 2D and 3D in vitro follicle culture [17–19], surgical intervention is required to obtain ovarian tissue and harvest cells. Moreover, their abilities to phenotypically-mirror theca cells upon co-culture with follicles has not been shown to date. Alternative cell sources, particularly those of renewable capacity (i.e. stem cells) present a unique advantage to generating cells that could give rise to a theca cell-like phenotype. To that end, stepwise in vitro differentiation of pluripotent stem cells into gonadal-like cells has been demonstrated in humans and mice [20–22] thus highlighting the potential to create a substitute cell source with prospective theca cell characteristics. Cell fate decisions that result in formation of the bipotential gonad in the embryo are the culmination of precisely timed cell signaling via gradients of morphogens such as FGF, BMP, and WNT proteins. Hence, protocols have been devised employing a stepwise induction of pluripotent stem cells using FGF, BMP, and WNT signaling such that cells progress through a mesoderm-like state, and finally a bipotential gonad-like state [23–25].

In this study, we describe the use of a PEG hydrogel culture system with or without native ovarian cells for the in vitro development of bovine preantral follicles and explore the potential of bovine embryonic stem cells to be induced towards gonadal-like cells in vitro. We first hypothesized that culture in PEG hydrogels would improve the growth and viability of bovine preantral follicles. By examining the gene expression of ECM-degrading enzymes of primary- and early secondary-stage follicles, we tailored the hydrogel design to respond to bovine follicle enzyme secretion. Based on the evidence that cell supplementation improves mouse follicle development, we also examined the effect of dissociated bovine ovarian cells (BOCs) on bovine preantral follicle growth and if BOCs exhibit a pre-theca cell phenotype when cultured in vitro using PEG hydrogels. Finally, we hypothesized that bovine embryonic stem cells (bESCs) can be differentiated into the intermediate mesoderm and early coelomic epithelium/bipotential gonad.

## MATERIAL AND METHODS

All materials were purchased from Thermo Fisher Scientific unless otherwise specified.

### Follicle Isolation and bovine ovary single-cell dissociation

Fresh bovine preantral follicles were isolated as previously described [26]. Follicles were maintained in warmed follicle wash media until subsequent experiments. For bovine ovary single-cell dissociation (n = 3 replicates), cortical tissue (250-300 mg) was fragmented into 500 µm^2^ pieces incubated with Hank’s balanced salt solution (Ca^2+^/Mg^2+^) containing 1 mg/mL collagenase IV, 1 U/mL DNase I, and 50 μg/mL Liberase (5401119001, Sigma Aldrich). The solution with tissue was shaken at 38.5 °C for 20 min, pipetted 15 times using a serological pipette, and shaken again for 20 min to mechanically aid in the dissociation of cells. Enzymatic activity was stopped by adding 20% v/v fetal bovine serum to the solution. The tissue-dissociated solution was filtered through a 100 μm then through a 35 μm cell strainer. Cells collected after final filtration were centrifuged (300 x*g* for 5 min) and the resulting pellet was resuspended in 1 mL Hank’s balanced salt solution without Ca^2+^/Mg^+^ until encapsulation.

### PEG hydrogel materials, preparation, and follicle encapsulation

PEG hydrogel experiments were conducted using 8-arm PEG vinyl sulfone (PEG- VS) (40 kDa, >99% purity, JenKem Technology) and crosslinked with MMP- and Plasmin-sensitive peptide sequences (Ac-GCRD**VPMS**MRGGDRCG**YKNS**CG, i.e. YKNS/VPMS) (2391.8 g/mol, >90%, Celtek Peptides). PEG and crosslinker peptide were dissolved in isotonic 50 mM HEPES buffer and mixed to a final composition containing 5% PEG-VS with 1:1 stoichiometric ratio of VS to thiol (side chain present on cysteines in peptide sequence). Sixteen-µl hydrogels containing preantral follicles were formed between parafilm-lined glass slides and allowed to gelatinize for up to 10 min. Then, hydrogels were transferred to warmed follicle maintenance media to swell and placed in a humidified incubator (37°C). Freshly isolated follicles were maintained in warmed follicle wash media until encapsulation. In experiment 1, follicles (n = 1-4) were first washed in 10% PEG-VS and either encapsulated in PEG hydrogels or placed into 2D culture. Hydrogel-encapsulated and 2D culture follicles were placed into individual wells of a 96-well plate containing follicle culture media (1:1 ratio of α-MEM and F12 with glutamine supplemented with 3 mg/mL bovine serum albumin, 100 ng/mL penicillin/streptomycin, 1 mM sodium pyruvate, 1 % (v/v) non-essential amino acids, 10 mg/l insulin, 5.5 mg/l transferrin and 6.7 μg/L selenium (ITS), 100 ng/mL human recombinant follicle stimulating hormone (228-12609-2, RayBiotech), and 50 μg/mL ascorbic acid. Half of the spent media was changed every 2 days and follicles were cultured for 10 days. To assess follicle growth over the culture period, the average of 2 perpendicular diameter measurements were taken at day 0, day 5, and day 10 using an inverted microscope (Revolve, Echo Labs). The percentage of growing and non-growing follicles (growth defined as having at least 5% increase in diameter by day 10 compared to day 0) and percent increase in size in growing-only follicles was calculated. In total, 15 replicates were conducted with a total of 216 follicles (n = 153 in PEG and n = 63 in 2D control)

### Cell viability staining and quantification

Bovine ovarian cells and MeLCs were resuspended in PEG-VS + crosslinker peptide VPMS/YKNS at a concentration of 1, 2, 10, and 20 x 10^6^ cells/mL to test cell viability immediately after encapsulation, 5 and 10 days after in vitro culture. Hydrogels were cultured individually in 96-well plates with 150 μl of follicle culture media with half media changes every 2 days. After culture, hydrogels were incubated for 10 min with pre-equilibrated alpha-MEM containing 2 μM propidium iodide (PI) and 1:1000 Hoechst (1 μg/mL final concentration). Negative control hydrogels were void of propidium iodide and used for setting exposure limit during imaging. Hydrogels were imaged using an epifluorescent microscope (Revolve) and the total number of cells and number of viable cells were quantified using ImageJ after converting to binary images where PI/Hoechst-positive cells were white, and background was black. Total number of cells positive for PI were divided by total cells quantified by Hoechst staining to determine percentage of unviable cells. Percentage of viable cells was determined by subtracting percentage of unviable cells from 100. The experiment was repeated 4 times using 3 hydrogels per cell density for each replicate.

### Bovine ovarian cells with follicle encapsulation and culture in PEG

To test the impact of BOC and co-encapsulation on follicle viability and growth and theca cell differentiation during in vitro culture, hydrogels were formed with either 1) only follicles, 2) 5 x 10^6^ BOCs/mL only, or 3) follicles and 5 x 10^6^ BOCs/mL BOCs. Each hydrogel contained 1-5 follicles (n = 65 follicles); 5 replicates). Hydrogel formation and follicle encapsulation was performed as previously described with the addition of staining follicles with the cell membrane marker PKH26 (2 μM final concentration; Sigma-Aldrich) prior to encapsulation to improve visualization. Hydrogels were cultured for 10 days with half of the medium changed every other day. Follicles were considered viable if the basement membrane was attached, the follicles were void of vacuoles, and granulosa cells were not darkened or mishappened. To recover cells for RT-qPCR, the hydrogels were incubated in phenol red-free alpha-MEM with Liberase (1.3 wünsch units/mL) and incubated at 37 °C and 5% CO_2_ for 30 min with repeated pipetting to degrade the hydrogel and extract the cells. After centrifugation, the cell pellet was used for RNA extraction, cDNA synthesis and RT-qPCR as described below.

### Culture of bovine embryonic stem cells (bESCs)

Bovine embryonic stem cells (n = 2 female lines) were routinely cultured feeder-free in NBFR medium (N2B27 medium [1:1 DMEM/F12 and Neurobasal media, 0.5% v/v N-2 supplement, 1% v/v B-27 supplement, 2 mM MEM Non-Essential Amino Acid solution, 1% v/v GlutaMAX supplement, 0.1 mM 2-mercaptoethanol, 100 U/mL Penicillin, and 100 μg/mL Streptomycin] supplemented with 1% bovine serum albumin (BSA), 20 ng/mL bFGF (100-18B, PeproTech), 2.5 μM IWR-1 (I0161, Sigma), and 20 ng/mL Activin A (338-AC, R&D Systems)]. Cells were passaged onto vitronectin-coated plates every 2-4 days using TrypleExpress at 1:4-1:6 split ratio and cultured in humidified incubators at 37 °C and 5% CO_2_. When cells were passaged, media was supplemented with 10 μm ROCK inhibitor Y-27632. Media was changed daily and bESCs used for mesoderm induction were between passage 18-26.

For mesoderm induction, bESCs were passaged onto 12-well vitronectin-coated plates at 40,000 cells per well in mesoderm-induction media [GMEM medium, 15% knockout serum replacement, 0.1 mM NEAA, 2mM GlutaMAX, 1mM sodium pyruvate, 0.1mM β-mercaptoethanol, 100 U/mL Penicillin, and 100 μg/mL streptomycin and supplemented with 3 μm CHIR99021 and 70 ng/mL Activin A]. Cells were cultured in mesoderm induction medium for 48 hours with daily medium change and thereafter the cells were considered “mesoderm-like cells” (MeLCs) and used for subsequent experiments.

Mesoderm-like cells were cultured in mesoderm induction medium without activin A for an additional 72 hours and supplemented with 10 ng/mL bFGF beginning immediately after MeLC induction, 24 hours after MeLC induction, 48 hours after MeLC induction, or no supplementation (mesoderm medium only). Based on the finding of which bFGF regimen best upregulated gene expression, in a second experiment, MeLCs were cultured in mesoderm induction media without activin A, with 10 ng/mL bFGF and with either 0, 1, 10, or 20 ng/mL of BMP4 for an additional 72 hours. All experiments contained duplicate wells for each condition tested and had 3 biological replicates/independent differentiations. Cells were imaged every day during each differentiation on the EVOS (Thermo Fisher) inverted microscope.

### End-point (RT-PCR) and quantitative reverse transcription PCR (qRT-PCR)

Freshly isolated follicles were pooled by stage and snap frozen in minimal PBS. To examine the expression of transcripts for extracellular-matrix degrading enzymes, primary follicles (n = 3 pools of 46-56 follicles per pool) and early secondary follicles (n = 3 pools of 34-40 follicles per pool) were subjected to RNA isolation using a protocol with a combination of Trizol, Purelink DNase treatment, and Qiagen RNeasy Micro Kit. Final total RNA was eluted into 14 μl of RNase-free water. Bovine ovarian cells (BOCs), bESCs, MeLCs and differentiated cells were pelleted, snap frozen, and subjected to RNA isolation following manual instructions using the Qiagen RNeasy Micro Kit. Maximum available RNA mass was used for cDNA synthesis of primary and early secondary follicles and 250 ng - 1 μg of total RNA was used for cDNA synthesis of bESCs, BOCs, MeLCs, and differentiated cells (RevertAid cDNA Reverse Transcription Kit). For RT-qPCR, 1-2 μl of cDNA (50 ng total), 1x SsoAdvanced Universal SYBR Green Supermix (Bio-Rad, Hercules, CA), and 250 nM of forward and reverse primers were mixed for 10 µl total volume and pipetted into 96-well plates. The reaction was carried out in the CFX96 Touch Deep Well Real-Time PCR Detection System (Bio-Rad). Cycling conditions were an initial denaturation step at 95 °C for 30 sec followed by 40 cycles of denaturation at 95 °C for 10 sec, annealing at 60 °C for 30 sec, and extension at 60 °C for 5 sec. To confirm specificity of PCR products for each gene, each assay included a melt curve analysis and non-template control. Data was normalized to *ACTB* by the delta-Ct method and relative expression data are represented as 1/ΔCT. For conventional PCR, PCR products from quantitative PCR reactions were ran on a 2% agarose gel to visualize the PCR products. For RT-qPCR experiments using *BAX* and *BCL2* primers, a standard curve of fresh bovine ovarian cells was used to estimate absolute quantity of mRNA present in unknown samples of BOCs. The ratio of *BAX*/*BCL2* was calculated from mRNA quantity of respective genes and used to determine if cells were in a pro- or anti-apoptotic state. All primers used in these experiments are listed in Table 1.

**Table 1.**
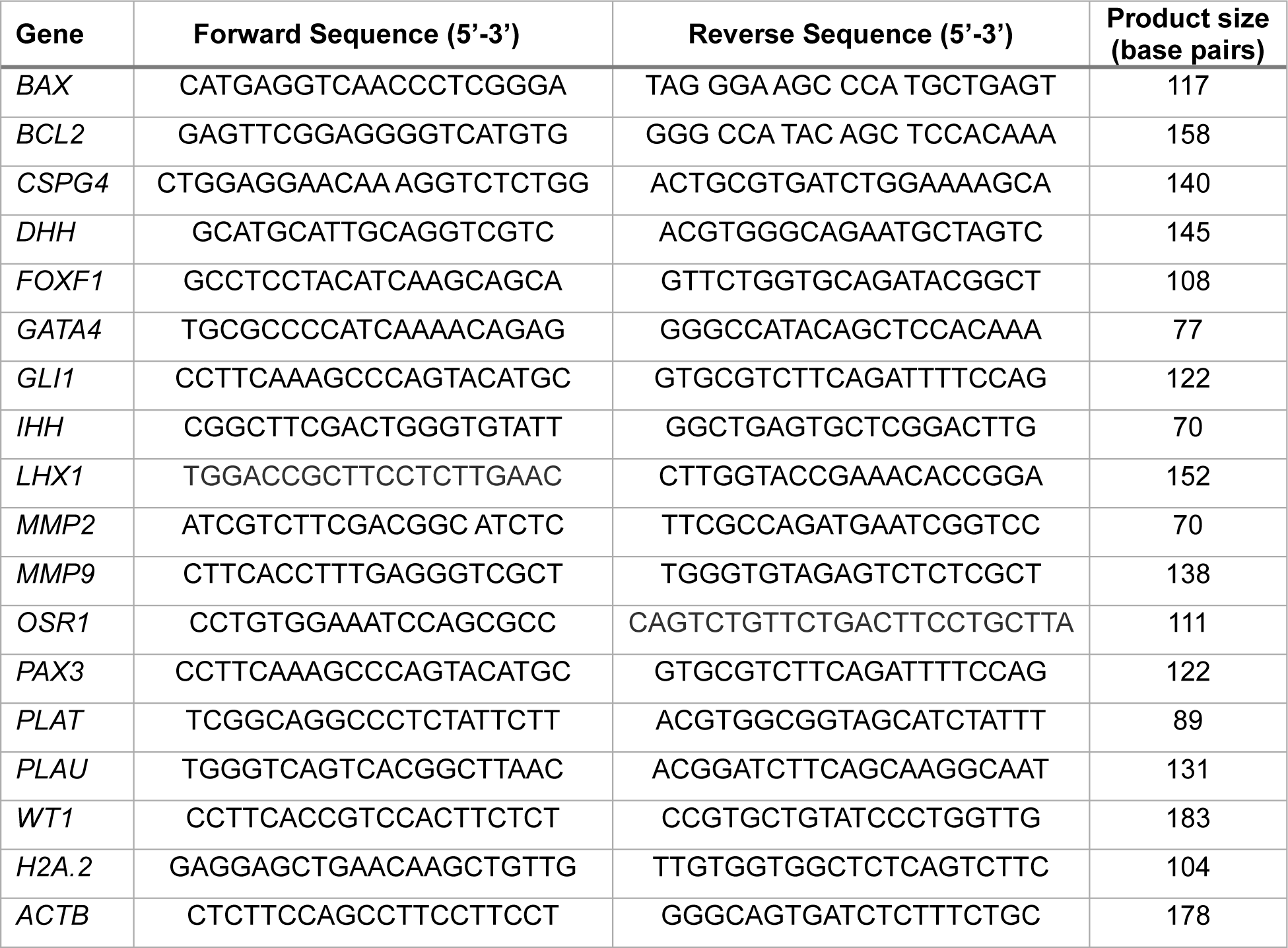
Primer list.

### Statistical analysis

All data was first assessed for normality using a Shapiro-Wilk test. PEG growth data was subjected to a two-way ANOVA using RStudio with effect of replicate included in the model. Cell viability and RT-qPCR data were subjected to a one-way ANOVA or Kruskal-Wallis test if data was not normally distributed using Graph Prism. Post-hoc tests (Tukey’s and Dunn’s for ANOVA and Kruskal-Wallis, respectively) were conducted when appropriate. RT-qPCR data were analyzed based on delta-Ct values. Comparisons were considered significantly different when associated with *P* < 0.05.

## RESULTS

To design a proteolytically-degradable PEG hydrogel that would most likely be compatible with bovine preantral follicles, we first examined the gene expression of ECM-degrading enzymes in primary and early secondary follicles that express the key follicle marker *FSHR*. Matrix metalloproteinases (MMPs) and plasminogen activators are ECM-degrading enzymes and play a pivotal role in tissue remodeling. We found the bovine primary and early secondary follicles express *MMP2*, although in one pool we only found *MMP2* in early secondary follicles. However, neither follicle stage expressed *MMP9* (Figure 1A). Plasminogen-activator urokinase (*PLAU*) was expressed in primary and early secondary follicles from one pool, but not in the second or third pool. Similarly, Plasminogen-activator tissue (*PLAT*) was expressed in primary follicles of one pool and in early secondary follicles from two other pools (Figure 1A). Since we collectively found mRNA expression of *MMP2* and plasminogen enzymes, we proceeded with using a PEG hydrogel that is functionalized with VPMS (MMP-sensitive) and YKNS (plasminogen-sensitive) peptide sequences (Figure 1B).

**Fig 1.**
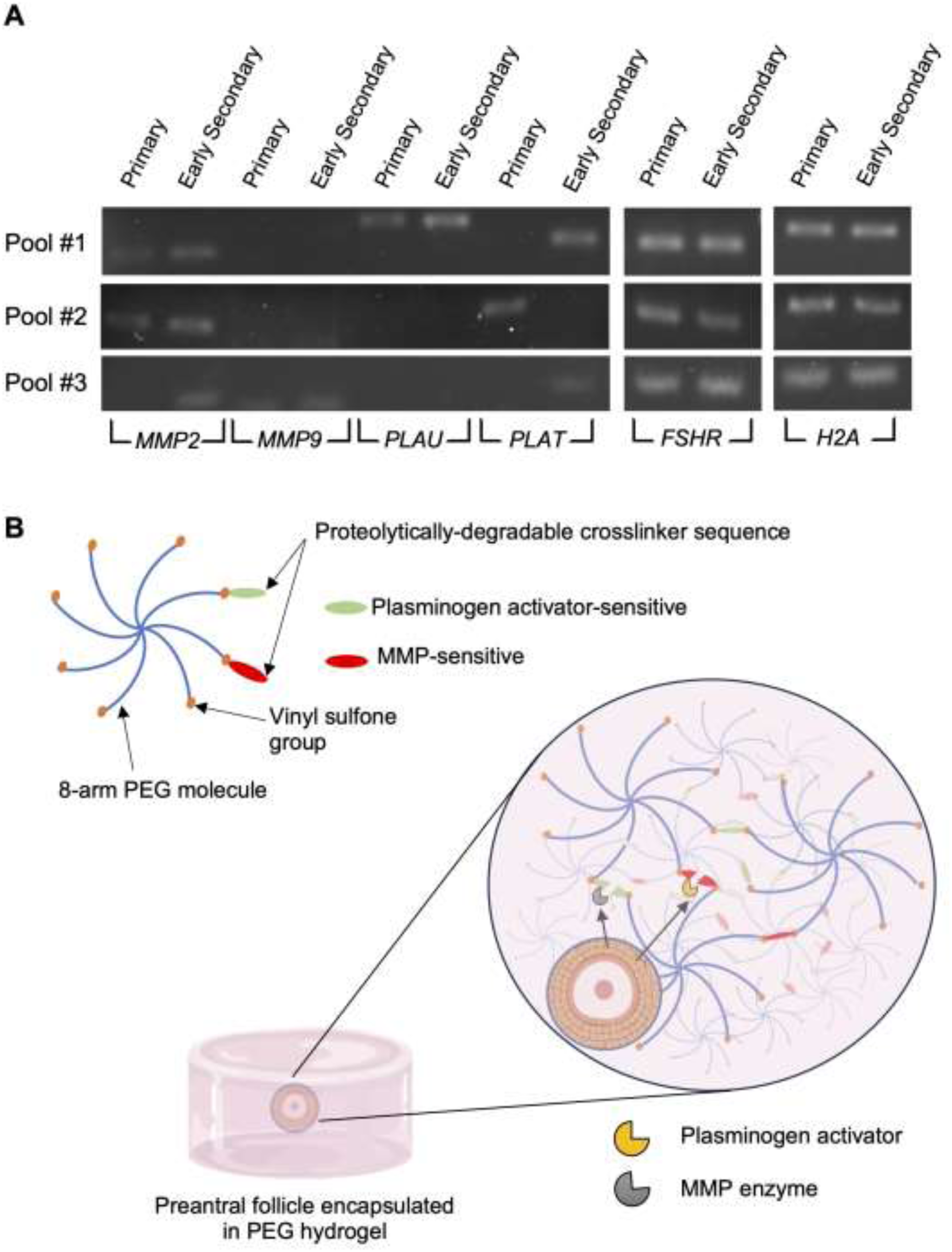
Bovine preantral follicles express extracellular matrix-degrading enzymes, allowing the design of a PEG hydrogel containing target peptides for controlled degradation. (**A**) Agarose gel with products of RT-PCR of primary and early secondary follicles expressing mRNA for *MMP2, MMP9, PLAU, PLAT, FSHR,* and *H2A.* Each pool of primary and early secondary follicles (n = 3 pools per stage) contained 46-56 and 34-40 follicles, respectively. (**B**) Schematic of PEG hydrogel for follicle encapsulation. PEG = poly(ethylene glycol). MMP = matrix metalloproteinase

Next, we tested the hypothesis that bovine preantral follicles are more likely to grow and demonstrate greater growth rate when encapsulated in PEG hydrogels compared to 2D traditional methods after 10 days of culture. We found no difference in the percentage of growing follicles between PEG (35.8% ± 6.23%) and 2D control (30.3% ± 7.88%) (*P* = 0.56) (Figure 2A). We found that replicate (i.e. individual ovary) affected the percentage of growing follicles (*P* < 0.05) regardless of culture condition. It is worth noting that in the control group, no follicle growth was observed in four out of the 15 replicates, whereas in the PEG group there was growth, although variable, in every replicate (Figure 1A). Of the replicates where at least one follicle demonstrated growth in either condition, we found no difference (*P* = 0.50) in the growth rate from initial size (PEG: 18.8% ± 3.32%, control: 22.7% ± 3.49%) (Figure 2B). There was no effect of replicate on growth rate (*P* = 0.78).

**Fig 2.**
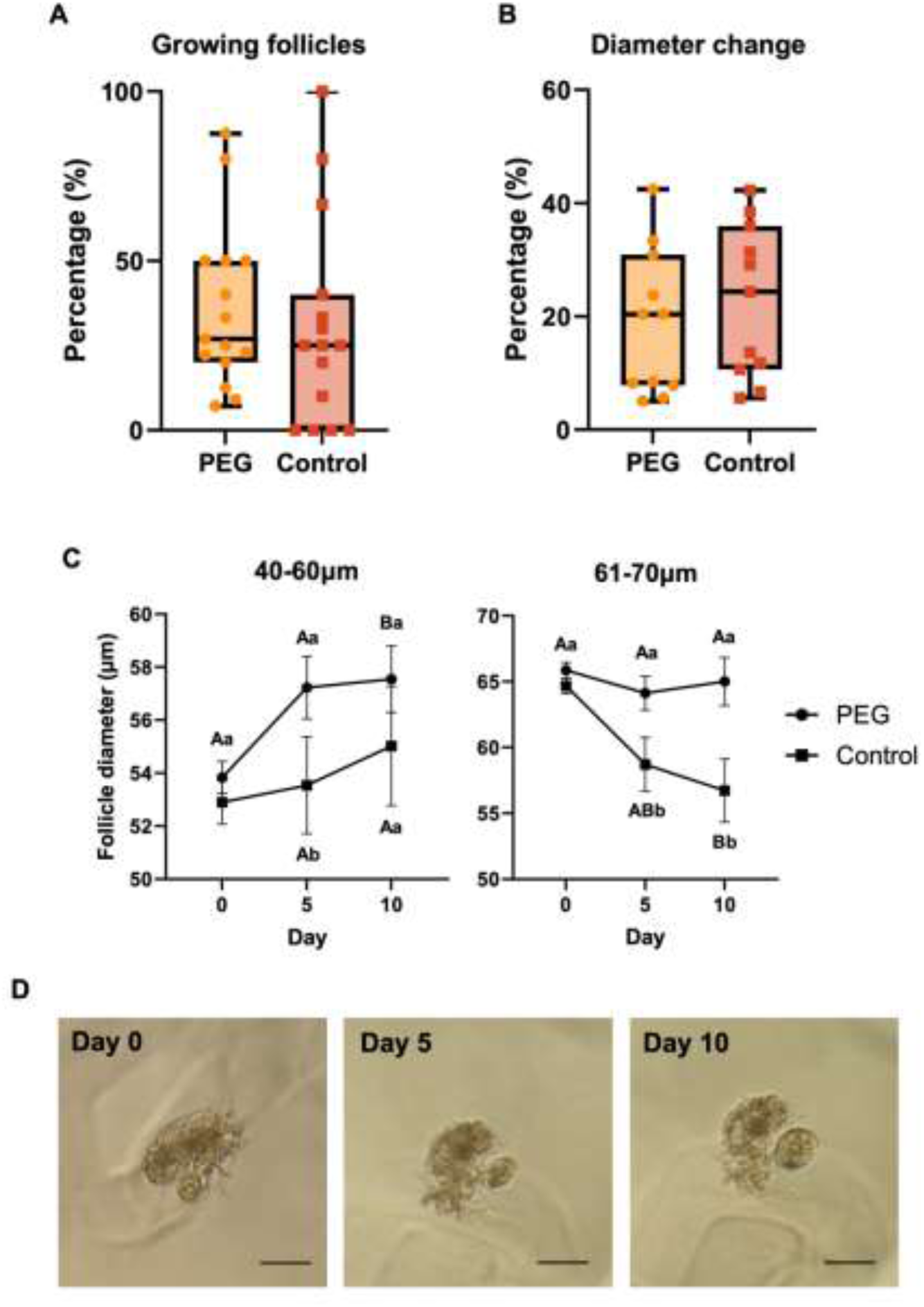
Bovine preantral follicle in vitro growth using PEG or 2D control culture systems. Percentage of preantral follicles that grew (**A**) and percent change in diameter from follicles that grew (**B**). (**C**) Bovine preantral follicle diameter growth during in vitro culture at indicated starting diameters/stages. Lowercases letters that differ = significant difference (*P* < 0.05) between treatment groups in the same day. Uppercase letters that differ = significant difference (*P* < 0.05) within the same treatment group but compared to day 0. Data are presented as mean ± SEM. (**D**) Representative images of follicle grown in PEG over 10-day culture period. Scale bar = 100 µm.

We further examined the diameter changes between day 0, 5, and 10 in follicles within a narrow range of starting diameters corresponding to their stage of development between the primary to early secondary. When we examined follicles with starting diameters corresponding to the early primary stage (40-60 μm), we found PEG hydrogels outperformed the control conditions as day 5 follicles were larger in PEG compared to 2D control (*P* < 0.05). Moreover, by day 10, PEG hydrogel follicles had a significant increase in diameter compared to day 0 (*P* < 0.05), whereas control follicles showed no difference (*P* = 0.72). Follicles starting at 61-70 μm showed a decrease in diameter by day 10 of culture in control conditions (*P* < 0.05), whereas follicles cultured in PEG maintained a similar diameter over time (*P* = 0.68; Figure 2C). Therefore, day 5 and 10 follicles were significantly smaller (*P* < 0.05) in control follicles when compared to similar timepoints of PEG follicles. A representative image of a follicle grown in a PEG hydrogel over the 10-day period is depicted in Fig. 2D.

We evaluated cell viability of BOCs when encapsulated in PEG hydrogels and cultured for 10 days (Figure 3A-C). When encapsulated at 1 or 2 x 10^6^ cells/mL, BOCs showed low cell viability (data not shown), whereas concentrations of 10 or 20 x 10^6^ cells/mL showed high viability (70-90%) and cell aggregation (Figure 3B and C). In this experiment, we found that neither control (no BOCs encapsulated) or hydrogels with BOCs supported follicle growth over the 10 day-culture period (Figure 3D). Moreover, gene expression analysis of pre-theca cell markers in the BOCs after 10 days of culture revealed a decrease in expression of *PTCH1* (*P* < 0.01) and *GLI1* (*P* < 0.01) when compared to day 0 fresh cells in both BOCs cultured alone and co-encapsulated with follicles (Figure 3E). We found a decrease in both *BAX* and *BCL2* in BOCs that were cultured with follicles and a decrease in *BCL2* in BOCs cultured in PEG hydrogels alone (*P* < 0.05; Figure 3F). However, there was no difference in *BAX*/*BCL2* ratio, thus indicating no difference in the pro-versus anti-apoptotic balance between cultured and fresh BOCs. Given that native BOCs did not show maintenance of pre-theca identity or contribute to follicle growth, we then explored the differentiation of bESCs into progenitor cells of the bipotential gonad by testing effect of WNT, bFGF and BMP4 signaling (Figure 4A). We first tested the hypothesis that MeLCs exposed to a pulse of bFGF during intermediate mesoderm/bipotential gonad differentiation would upregulate gene expression of key markers at the end of induction (Figure 4B and C). Compared to bESCs and 0, 24, 48, and 72 hours of 10 ng/mL bFGF exposure, MeLCs had higher expression of the early lateral plate/intermediate mesoderm marker *LHX1* (Figure 4B; *P* < 0.001). We also found the intermediate mesoderm marker *OSR1* to be upregulated in all bFGF regimens compared to bESCs (*P* < 0.01). When analyzing expression of bipotential gonad markers, there was no difference in *GATA4* (*P* = 0.15) across all experimental groups. We also found higher expression of *WT1* after 0, 24, 48, and 96 hours of bFGF (i.e., after 4 additional days of culture) compared to bESCs and MeLCs (*P* < 0.001). There was no difference in expression of *WT1* between bESCs and MeLCs (*P* = 0.15). There was also no difference in expression of the paraxial mesoderm marker *PAX3* across all experimental groups (*P* = 0.06), indicating induction conditions did not show strong propensity to drive cells to the paraxial mesoderm fate. However, the lateral plate mesoderm marker *FOXF1* was significantly upregulated compared to bESCs in 0, 24, and 48 hours of bFGF exposure (*P* < 0.05).

**Fig 3.**
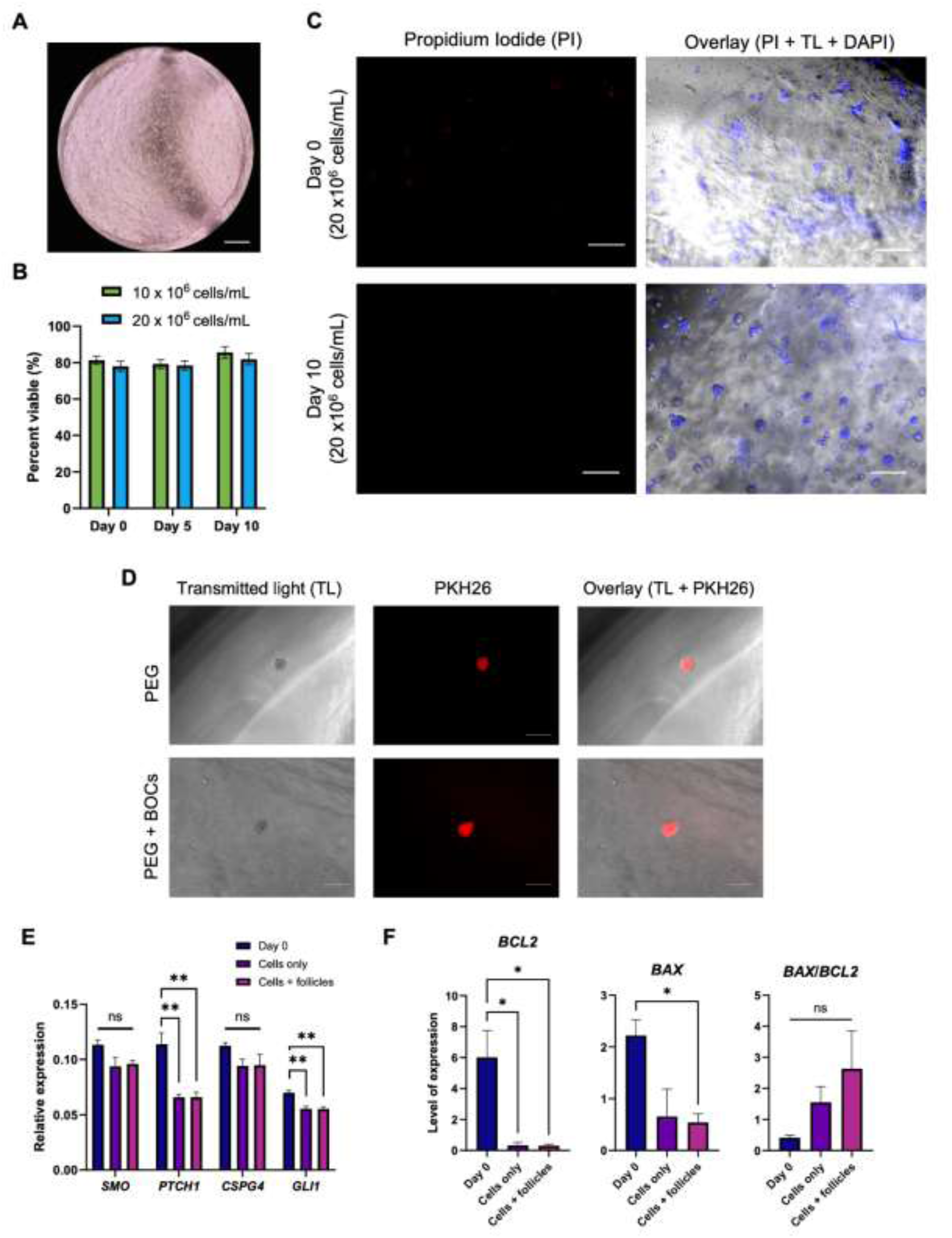
Co-culture of bovine preantral follicles with bovine ovarian cells (BOCs) in PEG hydrogels. (**A**) Representative image of BOCs in PEG hydrogel (scale bar = 500 µm). (**B**) Viability of BOCs encapsulated in PEG hydrogels during in vitro culture. (**C**) Propidium iodine (PI) staining in BOCs encapsulated in PEG hydrogels at day 0 and day 10 of culture. Counterstained with DAPI (TL = transluminescent; scale bar = 100 µm). (**D**) Unviable preantral follicle encapsulated in PEG hydrogel with or without BOCs using PKH26 to label follicles at day 10 of culture (scale bar = 100 µm). (**E**) Gene expression of pre-theca cell markers in BOCs (relative expression = 1/ΔCT). (**F**) Anti- and pro-apoptotic gene marker expression in BOCs (level of expression = absolute quantification from standard curve). Data are presented as mean ± SEM.

**Fig 4.**
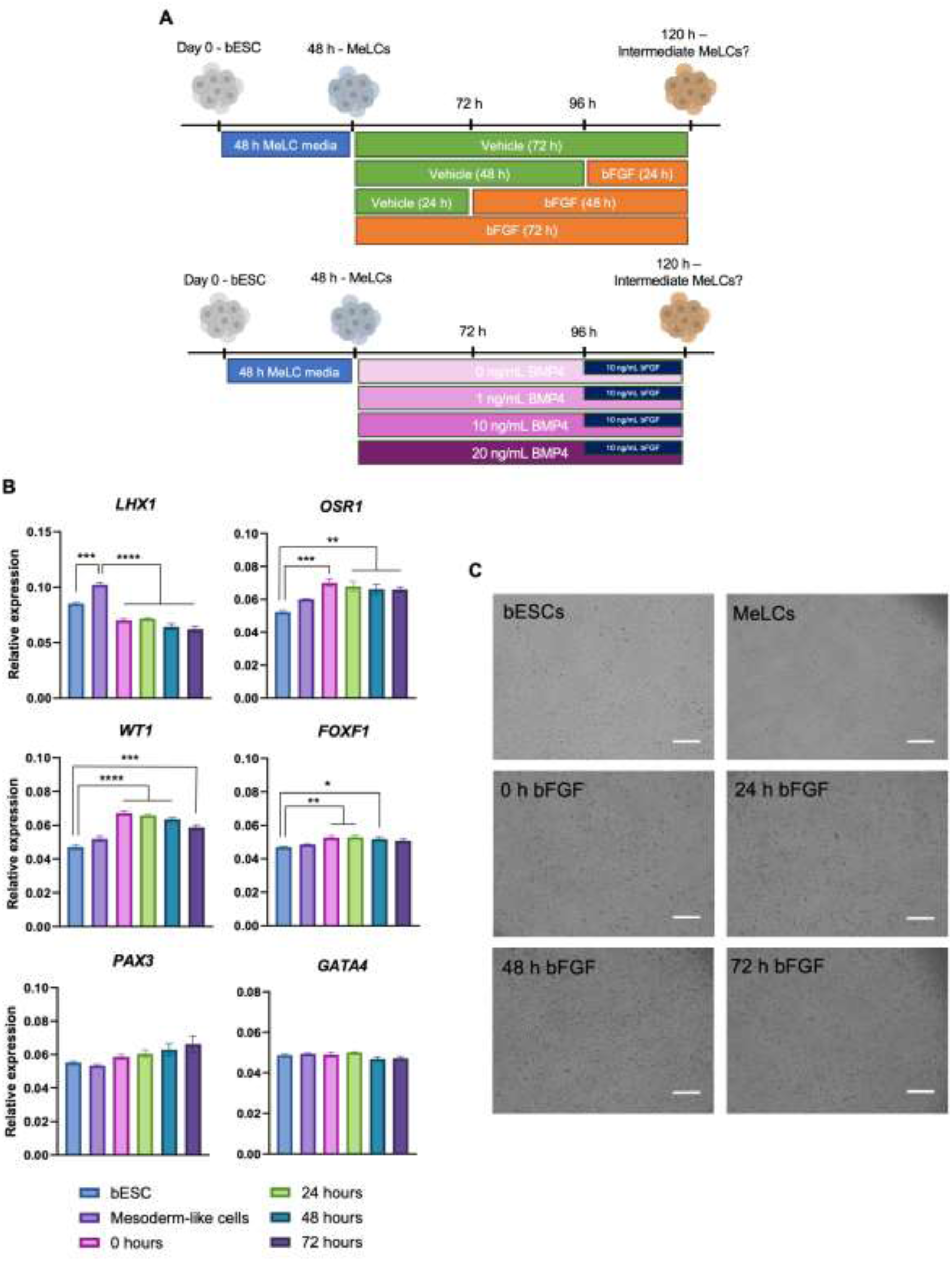
In vitro differentiation of bovine embryonic stem cells (bESCs) towards progenitors of somatic bipotential gonad-like cells. (**A**) Schematic diagram of experimental design for bFGF and BMP4 experiments. (**B**) Gene expression of intermediate mesoderm, bipotential gonad, paraxial and lateral plate mesoderm markers before and after bESC differentiation using bFGF at various days of exposure. (**C**) Representative images of cells from each treatment group. Data are presented as mean ± SEM. Significant differences are noted by * = *P* <0.05, ** = *P* <0.01, *** = *P* <0.001, **** = *P* <0.0001.

Because all bFGF regimens tested caused elevation in *WT1* and *OSR1* expression, we chose to use a 24-hour pulse of bFGF in the next experiments when testing the synergistic effects of including various concentrations of BMP4 over the same culture period time and in two bESC lines (Figure 5A and B). Similar to the bFGF experiment, we found the highest level of *LHX1* expression in MeLCs (Figure 5A; *P* < 0.0001). However, 10 and 20 ng/mL of BMP4 led to lower expression when compared to bESCs (*P <* 0.05). Also confirming the previous results, *OSR1* was elevated in all experimental groups, including MeLCs, when compared to bESCs (*P <* 0.01). Unlike in the bFGF experiment where there was no difference amongst exposure times to bFGF, we found an increase in *GATA4* in MeLCs and when 1, 10 or 20 ng/mL of BMP4 were present (*P* < 0.05). *WT1* was upregulated in 0, 1, 10, or 20 ng/mL BMP4, but not in MeLCs (*P* < 0.05). Again, we found no difference in *PAX3* expression among all concentrations of BMP4 when compared to bESCs. As expected, increasing the concentration of BMP4 led to a significant increase in *FOXF1* (*P* < 0.05), indicating a likely shift towards the lateral plate mesoderm lineage with rising levels of BMP4 which is similar to in vivo development. We further assessed WT1 and OCT4 expression in bESCs, MeLCs, and cells cultured with 0 and 1 ng/mL BMP4 as these concentrations led to elevated *WT1* without greatly upregulating *FOXF1* (Figure 5B). We found positive nuclear staining for OCT4 in bESCs and MeLCs and no signal in cells exposed to 0 or 1 ng/mL BMP4, indicating likely loss of pluripotency after mesoderm induction. We found nuclear localization of WT1 in bESCs and nuclear and cytoplasmic localization in MeLCs (48 h culture) and 0 ng/mL BMP4 (120 h culture). Interestingly, we did not observe WT1 expression at the protein level when cells were exposed to 1 ng/mL BMP4.

**Fig 5.**
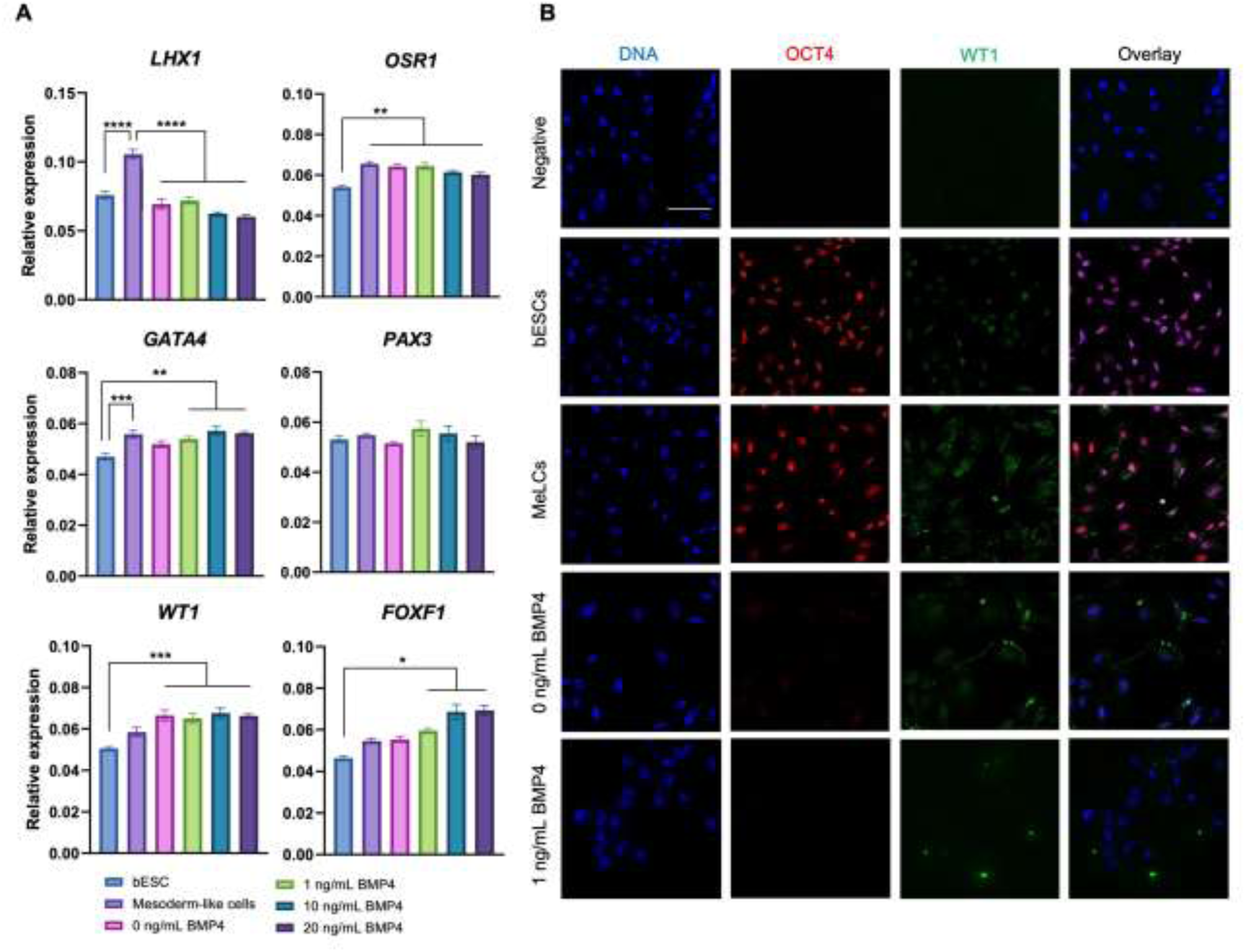
In vitro differentiation of bovine embryonic stem cells (bESCs) towards progenitors of somatic bipotential gonad-like cells testing BMP4 concentrations. (**A**) Gene expression of intermediate mesoderm, bipotential gonad, paraxial and lateral plate mesoderm markers before and after bESC differentiation (**B**) Representative images of cells from each treatment group. Data are presented as mean ± SEM. Significant differences are noted by * = *P* <0.05, ** = *P* <0.01, *** = *P* <0.001, **** = *P* <0.0001.

## DISCUSSION

To our knowledge, this study is the first to utilize a novel bioengineered PEG hydrogel system for in vitro culture of bovine preantral follicles and assess whether bESCs could be differentiated towards progenitor cells of the bipotential gonad using stepwise in vitro culture protocols. The ultimate goal of this work is to develop novel assisted reproductive technologies that enable the use of small preantral follicles in conjunction with renewable theca-like cells for oocyte production. Since maintaining structural integrity of the follicle is a main concern with 2D culture, hydrogels represent a viable option as they provide framework to maintain the 3D configuration of follicles while permitting growth. Indeed, the theory to build a culture system complete with extracellular matrix components like the ovary has been suggested when early studies first explored bovine preantral in vitro development [27]. Further adaptations to include sophisticated ECM components in the hydrogel design have permitted mouse follicle growth in vitro starting as early as the primordial stage [28]. Our finding that ECM-degrading enzymes such as MMPs, PLAT and PLAU are expressed by bovine preantral follicles agrees with previous investigations in follicles and spent medium [37,38]. However, the fact that these transcripts were not uniformly expressed in all pools examined could indicate an intrinsic variability in the ability of the follicles to remodel the ECM. This could explain at least in part the variability observed in growth rate in the present study. Although we found that PEG hydrogels outperformed 2D control over 10 days in primary follicles, there was a notable difference in the percentage of follicles that grew between replicates (each represented by one ovary), further underlining the biological variability between samples and possibly an inherent propensity of some follicles for growth versus atresia. A limitation of this study using slaughterhouse-derived oocytes is the inability to track factors known to affect follicle viability such as animal age, nutritional and health status. Of note, early secondary follicles showed a stagnant rate of growth compared to primary follicles cultured in PEG hydrogels; this stagnation suggests that either 1) the 10 day-culture conditions remain substandard for bovine preantral follicles or 2) follicles canonically slow their growth at this stage of development which is a phenomenon seen in mice [29]. It is worth noting that in vivo, 10 days represents a very narrow window within the estimated 180 days of folliculogenesis in the cow, and therefore relatively small changes in growth could be expected in normally-developing follicles.

We tested the hypothesis of enhancing the bovine follicle culture system with cellular supplementation as others have shown that the addition of ovarian stromal cells, fibroblasts, and adipose-derived stem cells promote survival and growth of preantral follicles in several species [13,17–19,30]. When BOCs were encapsulated in PEG hydrogels and cultured, we found cell aggregates which may be due to absence of basement membrane binding proteins in the PEG composition, rendering the cells unable to adhere to the matrix and resulting in self-aggregation. Indeed, cell-extracellular matrix interactions are important for mediating complex cellular responses such as migration, proliferation, and differentiation [31], thus BOCs were likely incapable of integrating well throughout the hydrogel. Furthermore, the finding of decreased expression of genes known to be important for the pre-theca cell phenotype after culture shows that the BOCs did not maintain their identity, which could have contributed to the lack of success of this 3D co-culture system. Others have reported that mouse ovarian stromal cells transition from a predominately theca-like identity to almost entirely macrophage identity after 12 days of in vitro culture with follicles, partially corroborating our findings of theca cell loss [32]. Since we found similarly high signs of atresia between follicles co-encapsulated and follicles in PEG-only hydrogels, we pose the possibility that the prolonged protocol required to dissociate ovarian tissue, isolate and label follicles, and process for encapsulation may have contribute to decrease follicle viability beginning at day 0. In future experiments, pre-theca cells could be sorted (pending availability of reliable antibodies against specific theca cell surface markers) before incorporation into the PEG hydrogel. Furthermore, culture conditions that specifically support theca cell phenotype would likely need to be investigated.

In an attempt to produce gonadal-like cells for supplementation as pre-theca-like cells in follicle culture, we employed bovine embryonic stem cells to recapitulate mesoderm formation and early differentiation of the bipotential gonad with defined media conditions. We show that inclusion of exogenous signaling factors such as the GSK3β inhibitor CHIR 99021 for activation of WNT. WNT signaling can drive bESCs into intermediate mesoderm-like cells that show early signs of coelomic epithelial-like phenotype as indicated by gene and protein expression. During embryonic development, the sexually dimorphic gonad (ovary or testis) must develop beginning from the mesoderm germ layer, and then further differentiate into the intermediate mesoderm, then coelomic epithelium which ingresses and gives rise to the sexually amorphic “bipotential gonad” [33,34]. More specifically, upon Nodal, WNT and FGF signaling to induce mesoderm differentiation [35], a subset of cells in the caudal aspect of the early gastrulated embryo are specified to become precursors of the bipotential gonad. Signaling by WNT, Nodal, and FGF during anterior-posterior (A-P) body axis extension, which occurs at peri-gastrulation and is regulated by TBXT and other morphogens in the mesoderm, drives cell fate decisions that result in caudal body axis formation [36,37]. Concomitantly, mediolateral segmentation occurs when a low, medium, and high gradient of BMP signaling drives mesoderm cells into the paraxial, intermediate, and lateral plate mesoderm domains, respectively [38,39]. Following this knowledge of developmental biology, we chose to modulate WNT, Nodal, FGF and BMP signaling to mimic the molecular events that drive cell differentiation for pre-gonad development during early embryogenesis as shown in several mammalian species [34]. However, through these experiments, we found that under WNT activation, bFGF and BMP4 are not essential to induce the intermediate mesoderm or early coelomic epithelium. We observed peak expression of the intermediate mesoderm marker *LHX1* after 48 h of mesoderm induction under WNT and activin A, the latter a known inducer of *Lim1* (i.e. *Lhx1*) that has an important role in IM specification/kidney development [40]. We also found *OSR1* upregulated in all conditions after MeLC induction. *OSR1* is an early marker of IM and others have shown its necessity for the urogenital phenotype [41–43] and generating somatic cells of the gonad from PSCs [44]. Thus, we propose these cells may be reaching the IM state with possible capability of becoming progenitor bipotential gonad-like. Nevertheless, *OSR1* is not exclusive of the IM or gonads; kidney progenitor and lateral plate mesoderm cells express *OSR1* [43,45] and this marker is used to demarcate kidney progenitor-like cells during in vitro differentiation of PSCs [46–48]. Therefore, our intermediate mesoderm/early coelomic epithelium-like cells should be further evaluated for markers of early nephrogenesis.

We additionally found a rise in *WT1,* which is also an early, and more restrictive, marker of gonadal progenitor cells [49], in induced cells across all bFGF and BMP4 regimens compared to bESCs. This indicates that continuous WNT activation for 3 additional days after initial mesoderm induction can provoke *WT1* expression. WT1 is found along the A-P axis of mice as early as E9.0, with overlapping expression of GATA4 and NR5A1 in the coelomic epithelium by E10.0 and onward [49]. Moreover, WT1 activates the NR5A1 promoter [50], thus driving early gonad development. Interestingly, *WT1* upregulation was accompanied by *GATA4* upregulation when 10 or 20 ng/mL of BMP4 was included in culture which aligns with other studies [51]. However, we found variable expression of the protein WT1 in cells cultured beyond MeLC induction and after 3 additional days under 0 or 1 ng/mL of BMP4. Although typically found in the nucleus, WT1 isoforms with capability of shuttling between the nucleus and cytoplasm, therefore we postulate various WT1 isoforms with nuclear and/or cytoplasmic activity may be expressed in our cells [52]. We additionally highlight that there was no difference in *PAX3* in all experiments, demonstrating that our culture conditions did not favor differentiation of the paraxial mesoderm lineage. Interestingly, increasing BMP4 concentrations led to increased expression of *FOXF1* (lateral plate mesoderm marker), which aligns with in vivo studies showing a higher gradient of BMPs specifies the lateral plate of developing embryos [39].

Our findings provide seminal context to previous work looking at differentiation of ESCs from humans and mouse to gonad-like cells by using combinations of BMP4, FGF, CHIR, and activin A [53–55]. Alternative approaches to inducing gonad-like cell formation include overexpression of key transcription factors such as *SF1*, *WT1*, *RUNX1*, and *GATA4* [56,57]. The efficiency of induction can be variable when using small molecules or genetic engineering for differentiating stem cells, therefore heterogeneity in cell identity would be expected in the present study. Although not done in this study, cell-sorting based on expression of key markers could aid in purifying cell populations of interest and used for downstream in vitro culture to achieve bipotential and even ovary or testis-like cells. Yet, in vivo early somatic progenitor cells of the bipotential gonad are not specified homogeneously, therefore some heterogeneity could be warranted, especially when forming organoids as self-organizing structures. Finally, future studies should examine early bovine embryonic development in vivo as findings from these studies could shed light on the species-specific molecular mechanisms that drive gonad formation and inform future in vitro culture conditions to recapitulate the process.

In conclusion, these findings provide a key step in expanding current understanding of in vitro preantral folliculogenesis using a non-rodent model as well as sheds light on methodologies in differentiating bESCs into precursor-like cells of the ovary. The convergence of these two technologies will be pivotal to broadening the knowledge base of follicle development and early gonadogenesis such that novel assisted reproductive technologies can be expanded.

## ACKNOWLEDGEMENTS

The authors acknowledge Stephanie McDonnell and Amanda Morton for aiding in follicle collection and isolation. Funding: NIH 1F31HD111173, UC Davis Jastro Shields Graduate Research Award, and Gary B. Anderson Food Animal Fellowship to JC; USDA-NIFA W4112, 4171 multi-state groups. Some figures created with BioRender.com.

## AUTHOR CONTRIBUTION

Conceptualization: Juliana Candelaria, Anna Denicol; Methodology: Juliana Candelaria, Carly Guiltinan, Ramon Botigelli, Ariella Shikanov, Anna Denicol; Formal analysis and investigation: Juliana Candelaria, Anna Denicol; Writing - original draft preparation: Juliana Candelaria; Writing - review and editing: Juliana Candelaria, Anna Denicol, Ariella Shikanov, Carly Guiltinan, Ramon Botigelli,; Funding acquisition: Juliana Candelaria, Anna Denicol; Resources: Anna Denicol, Ariella Shikanov; Supervision: Anna Denicol.

